# CompMap: an allele-specific expression read-counter based on competitive mapping

**DOI:** 10.1101/2021.02.12.431019

**Authors:** S. Sánchez-Ramírez, A. D. Cutter

**Affiliations:** Department of Ecology and Evolutionary Biology, University of Toronto, 25 Willcocks Street, Toronto ON, M5S 3B2, Canada

**Keywords:** RNA-seq, allele-specific expression, Python, simulations

## Abstract

**Summary:** Changes to regulatory sequences account for important phenotypic differences between species and populations. In heterozygote individuals, regulatory polymorphism typically manifests as allele-specific expression (ASE) of transcripts. ASE data from inter-species and inter-population hybrids, in conjunction with expression data from the parents, can be used to infer regulatory changes in *cis* and *trans* throughout the genome. Improper data handling, however, can create problems of mapping bias and excessive loss of information, which are prone to arise unintentionally from the cumbersome pipelines with multiple dependencies that are common among current methods. Here, we introduce a new, selfcontained method implemented in Python that generates allele-specific expression counts from genotype-specific map alignments. Rather than assessing individual SNPs, our approach sorts and counts reads within a given homologous region by comparing individual read-mapping statistics from each parental alignment. Reads that are aligned ambiguously to both references are resolved proportionally to the allele-specific matching read counts or statistically using a binomial distribution. Using simulations, we show `CompMap` has low error rates in assessing regulatory divergence.

**Availability:** The Python code with examples and installation instructions is available on the GitHub repository https://github.com/santiagosnchez/CompMap

**Supplementary information:** 

## 1. INTRODUCTION

Geneticists’ long-standing quest to understand functional variation within genomes has led to the appreciation that changes in regulatory regions are crucial in fine-tuning developmental controls of gene expression to influence phenotypes, in addition to effects of structural changes in protein-coding genes (The ENCODE Project Consortium, 2012; Signor and Nuzhdin, 2018). Non-coding mutations to *cis*-regulatory elements that are localized close to a gene and *trans*-regulatory factors that are encoded at a distant location in the genome provide the genetic variation for expression of a focal gene, which, in turn, can drive adaptive divergence between populations and between species (Jones *et al.*, 2012; Wray, 2007). Identifying genetic variants with *cis*-acting or *trans*-acting regulatory effects, however, presents a serious challenge, being difficult to disentangle without explicit allele-specific expression (ASE) information (Signor and Nuzhdin, 2018).

Current ASE approaches include lengthy pipelines with multiple dependencies aimed at within-species datasets (Rozowsky *et al.*, 2011), tissue-specificity (Pirinen *et al.*, 2015), and cross-population samples (Fan *et al.*, 2020), usually requiring reads to be mapped to a single genotype-specific reference, followed by variant-calling and phasing. Here, we present `CompMap`, a self-contained, variant-calling-free Python tool for read-counting ASE datasets. The motivation for developing `CompMap` stems from the need to have a pipeline-free tool that avoids limitations from variant-calling and generates read-counts compatible with other downstream read-counting software.

## 2. METHODS

### 2.2 Implementation

In brief, `CompMap` parses two F_1_ hybrid BAM files with RNA-seq reads mapped to each parental genome simultaneously and compares the alignment quality of each read against both references. Allele-specific reads are then competitively assigned, based on their relative mapping scores, and counted. `CompMap` is implemented in Python-3 relying on the *pysam* API library to read and parse BAM files and *numpy* (e.g., for the binomial distribution). It requires these basic input files:

- BAM files with F_1_ hybrid (offspring) RNA-seq data mapped to each parental genotype
- BED files with genomic coordinates of the genes of interest; one per parental genotype

Preferably, BAM files are sorted and indexed. The 4th column of BED files should include gene name and match for homologous regions between parental genomes (i.e., same name for both BED files). The `-h` or `--help` argument will print descriptions and other information to screen.

The two most important arguments are `--AS_tag` and `--NM_tag`, which indicate labels for the alignment score and number of mismatches tags, respectively. By default, `CompMap` will use tags from the `STAR` aligner (Dobin *et al.*, 2012). The user can specify read-tag labels of different aligners, such as `BWA` (Li and Durbin, 2010) and `Bowtie2` (Langmead and Salzberg, 2012). The `--star` argument will apply the `NH` tag when looking for multiple read matches, which may improve speed. Finally, users can specify the `--binom` tag to assign ambiguous read-counts probabilistically to the alternate parental alleles. The binomial probability *p* describes the fraction of non-ambiguous reads with better matches to one parental genome for the corresponding gene (1-p better matching the other parental genome) and *size* defines the total number of ambiguously matching reads. The default behavior deterministically allocates reads with proportions *p* and *1-p* without using binomial sampling.

We validated `CompMap` read-counting by comparing it to `featureCounts` (Liao *et al.*, 2014) and `HTseq-counts` (Anders *et al.*, 2015), replicating results of `featurecounts`. We therefore recommend using `featureCounts` for read-counting of parental RNA-seq data used in combination with `CompMap`.

### 2.3 Simulations

Gene length and sequence differences between alleles can potentially impact the power to accurately detect and quantify ASE. To validate our approach, we simulated protein-coding sequence datasets of homologous alleles with synonymous site divergence. We neglected divergence at nonsynonymous sites because of their rarity in biological data of close relatives and to avoid assumptions about the strength and direction of selection. The presence of nonsynonymous differences in real data will make our power analysis conservative with respect to detecting and quantifying ASE.

The simulation procedure first drew 1000 random protein lengths from a Gamma distribution (scale=1000, shape=1.35; minimum length 300 aa). Non-stop codons were picked at random to form a “transcript”, all of which started with the Methionine coding “ATG” and ended with any of the three stop codons. Synonymous-site substitutions were imposed to create a divergent version of every “transcript”: each 4-fold, 3-fold, and 2-fold degenerate codon was allowed to mutate to a synonymous codon, selecting alternate codons with equal probability. Synonymous-site divergence used one of three rates: (1) high *d_s_*=0.1 substitutions/site, (2) moderate c/=0.01 substitutions/site, and (3) low *d_s_*=0.001 substitutions/site. `CompMap`’s GitHub repository contains code for this simulation procedure.

For each “allelic” transcript sequence, we then simulated RNA-seq reads using the R package `polyester` (Frazee *et al.,* 2015). Fold-changes in expression were randomly sampled from an exponential distribution with sign (up- or down-regulation) assigned randomly. These generated data defined “true” ASE counts for comparison to estimates derived from `CompMap`. Similarly, we generated RNA-seq datasets representing expression differences in each homozygous parent. Fold-change values between “alleles” and “parents” were drawn from independent distributions.

To validate ASE counts from `CompMap`, we combined allele-specific reads into a single FASTA file for each replicate and mapped them to each transcript reference using `BWA-MEM` (Li and Durbin, 2010). We then generated BED files with coordinates for each transcript. The resulting BAM and BED files were fed to `CompMap` to perform ASE counts using the `--NM_tag` with `NM`, providing the `BWA-specific` read tag for number of mismatches.

Standard read counts for “parental” and true “allele” read data were performed with `featureCounts` (Liao *et al.*, 2014). Raw read counts were then analyzed in R and differential expression analyses conducted with `DESeq2` (Love *et al.*, 2014). We followed McManus *et al.*, (2010) to statistically assign genes to different regulatory divergence categories (e.g., *cis*-only, trans-only, *cis-trans* compensatory, *cis x trans).*

## 3. RESULTS AND CONCLUSION

`CompMap` recovered high accuracy in allele-specific counts for all three simulated proteincoding datasets that spanned 100-fold range of divergence (Figure 1a). Count accuracy was highest for genes with high divergence between alleles (*d_s_*=0.1). In the more challenging case of low divergence (*d_s_*=0.001), `CompMap` slightly overestimated the number of reads for a given allele (by a factor of approximately 1.75), being most pronounced among genes with a lower magnitude of expression, as expected. However, this effect does not strongly perturb analyses of differential expression (Figure 1b). High-divergence alleles also show lowest variance in the ASE log_2_-fold-difference between simulated allele-specific reads counted by `featureCounts` and `CompMap` (Figure 1b). Despite the low divergence dataset yielding widest variability in estimated differential expression, the mean centered close to zero indicates little bias (Figure 1b).

**Figure 1.**
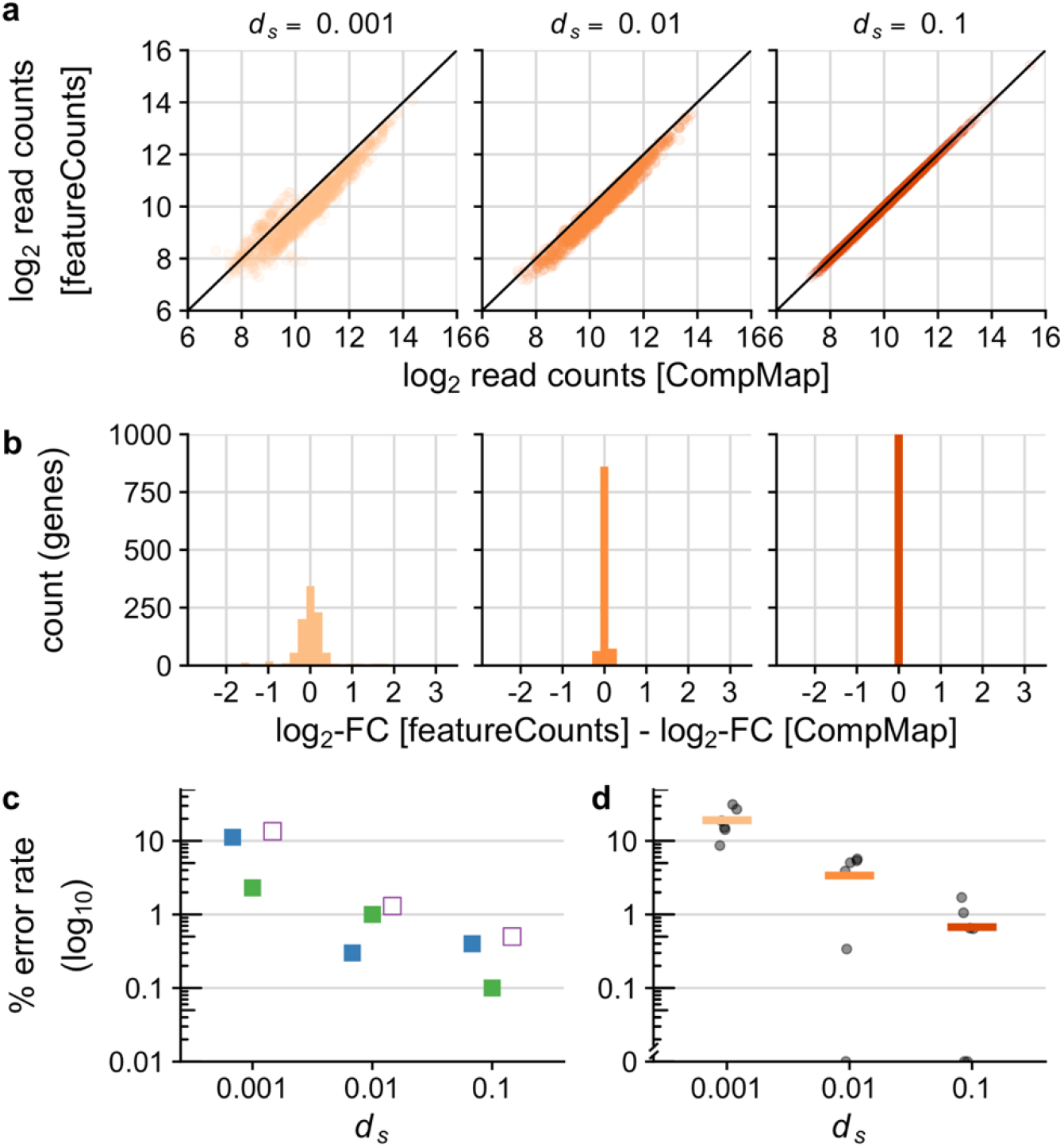
Accuracy in allele-specific expression (ASE) increases with protein-coding divergence using `CompMap`. a) Biplot of log_2_ raw read counts between “true” allele-specific counts using `featurecounts` and “mixed” read counts using `CompMap`. b) Histograms of the difference in log_2_-fold-change for ASE between `featureCounts` and `CompMap` (low *d_s_*=0.001: mean=0.0057, sd=0.508; moderate *d_s_*=0.01: mean=0.003, sd=0.068; high *d_s_*=0.1: mean=0.0017, sd=0.009). c) Estimates of the false negative error rate (blue), false positive error rate (green), and total error (open purple) to differential expression between alleles, and d) mean false negative rate across different regulatory divergence categories. See McManus *et al.*, (2010) for a complete list of regulatory divergence categories. The “y” axis on c and d is based on the percent error plotted on log_10_ scale.

Error rates to inferring differential expression between alleles declined approximately 10-fold with increasing divergence (Figure 1c). The power of `CompMap` to accurately classify genes into different regulatory divergence categories is greatest with high divergence between alleles (<2% false negative rates for the *d_s_*=0.1 dataset; <6% for intermediate *d_s_*=0.01) (Figure 1d). As expected, highest inaccuracy occurred with low divergence (*d_s_*=0.001 false negative rates ~20%).

Consequently, ASE reliability will be greatest for biological studies involving alleles with high sequence divergence, which will be challenging with neutral allele divergence ~0.1%, as within humans (Perry *et al.*, 2012) and *Caenorhabditis elegans* (Andersen *et al.*, 2012). Essential for such cases is a simulation framework for validation, as implemented in `CompMap` to assess the power to quantify ASE.

`CompMap`’s competitive read-mapping approach is sensitive and accurate, given sufficiently dense sequence differences between alleles. With `CompMap`, we 1) introduced a framework to reliably assess the power of recovering per-gene ASE read counts; 2) developed R code for simulation testing of coding sequence divergence; and 3) showed best ASE inference with ≥1% allelic synonymous-site divergence. `CompMap` is ideal for RNA-seq datasets derived from interspecies hybrids (Sánchez-Ramírez *et al.,* in press), as well as within-species analysis of systems with high genetic diversity, including *Drosophila* and *Caenorhabditis* (Cutter *et al.*, 2013).

## ACKNOWLEDGEMENTS

We thank the SciNet Consortium at the University of Toronto and Compute Canada for providing access to HPC resources.

## REFERENCES

Anders, S. et al. (2015) HTSeq-a Python framework to work with high-throughput sequencing data. Bioinformatics, 31, 166–169.

Andersen, E.C. et al. (2012) Chromosome-scale selective sweeps shape *Caenorhabditis elegans* genomic diversity. Nat. Genet., 44, 1–8.

Cutter, A.D. et al. (2013) Molecular hyperdiversity and evolution in very large populations. Mol. Ecol., 22, 2074–2095.

Dobin, A. et al. (2012) STAR: ultrafast universal RNA-seq aligner. Bioinformatics, 29, 15–21.

Fan, J. et al. (2020) ASEP: Gene-based detection of allele-specific expression across individuals in a population by RNA sequencing. PLoS Genet., 16, 1–23.

Frazee, A.C. et al. (2015) Polyester: Simulating RNA-seq datasets with differential transcript expression. Bioinformatics, 31, 2778–2784.

Jones, F.C. et al. (2012) The genomic basis of adaptive evolution in threespine sticklebacks. Nature, 484, 55–61.

Langmead, B. and Salzberg, S.L. (2012) Fast gapped-read alignment with Bowtie 2. Nat. Methods, 9, 357–359.

Li, H. and Durbin, R. (2010) Fast and accurate long-read alignment with Burrows-Wheeler transform. Bioinformatics, 26, 589–595.

Liao, Y. et al. (2014) featureCounts: an efficient general purpose program for assigning sequence reads to genomic features. Bioinformatics, 30, 923–930.

Love, M.I. et al. (2014) Moderated estimation of fold change and dispersion for RNA-seq data with DESeq2. Genome Biol., 15, 21–31.

McManus, C.J. et al. (2010) Regulatory divergence in *Drosophila* revealed by mRNA-seq. Genome Res., 20, 816–825.

Perry, G.H. et al. (2012) Comparative RNA sequencing reveals substantial genetic variation in endangered primates. Genome Res., 22, 602–610.

Pirinen, M. et al. (2015) Assessing allele-specific expression across multiple tissues from RNA-seq read data. Bioinformatics, 31, 2497–2504.

Rozowsky, J. et al. (2011) AlleleSeq: analysis of allele-specific expression and binding in a network framework. Mol. Syst. Biol., 7, 1–15.

Sánchez-Ramírez, S. and Cutter, A.D. (in press) Widespread misregulation of inter-species hybrid transcriptomes due to sex-specific and sex-chromosome regulatory evolution. PLoS Genet.

Signor, S.A. and Nuzhdin, S. V (2018) The evolution of gene expression in *cis* and *trans*. Trends Genet., 34, 532–544.

The ENCODE Project Consortium (2012) An integrated encyclopedia of DNA elements in the human genome. Nature, 488, 57–74.

Wray, G.A. (2007) The evolutionary significance of *cis*-regulatory mutations. Nat. Rev. Genet., 8, 206–216.

